# Exploring the utility of ssDNA aptamers directed against snake venom toxins as new therapeutics for tropical snakebite envenoming

**DOI:** 10.1101/2022.05.22.492967

**Authors:** Nessrin Alomran, Raja Chinnappan, Jaffer Alsolaiss, Nicholas R. Casewell, Mohammed Zourob

## Abstract

Snakebite is a neglected tropical disease that causes considerable death and disability in the tropical world. Although snakebite can cause a variety of pathologies in victims, haemotoxic effects are particularly common and are typically characterised by haemorrhage and/or venom-induced consumption coagulopathy. Antivenoms are the mainstay therapy for treating the toxic effects of snakebite, but despite saving thousands of lives annually, these therapies are associated with limited cross-snake species efficacy due to venom variation, which ultimately restricts their therapeutic utility to particular geographical regions. In this study, we sought to explore the potential of ssDNA aptamers as toxin-specific inhibitory alternatives to antibodies. As a proof of principle model, we selected snake venom serine protease toxins, which are responsible for contributing to venom-induced coagulopathy following snakebite envenoming, as our target. Using SELEX technology, we selected ssDNA aptamers against recombinantly expressed versions of the fibrinogenolytic SVSPs Ancrod from the venom of *Calloselasma rhodostoma* and Batroxobin from *Bothrops atrox*. From the resulting pool of specific ssDNA aptamers directed against each target, we identified candidates that exhibited low nanomolar binding affinities to their targets. Downstream ALISA, fibrinogenolysis, and coagulation profiling experiments demonstrated that the candidate aptamers were able to recognise native and recombinant SVSP toxins and inhibit toxin- and venom-induced prolongation of plasma clotting times and consumption of fibrinogen, with inhibitory potencies highly comparable to commercial polyvalent antivenoms. Our findings demonstrate that rationally selected toxin-specific aptamers can exhibit broad *in vitro* cross-reactivity against toxins found in different snake venoms and are capable of inhibiting toxins in pathologically relevant *in vitro* and *ex vivo* models of venom activity. These data highlight the potential utility of ssDNA aptamers as novel toxin-inhibiting therapeutics of value for tackling snakebite envenoming.

## Introduction

Snakebite envenoming is a significant public health issue as more than 5.4 million people are bitten annually by venomous snakes, and it mainly affects resource-poor communities in the rural tropics and subtropics across Africa, the Americas, Asia, the Middle East, and Australasia (1-3). Since 2017, snakebite has been classified by the World Health Organization as a neglected tropical disease (4). Snake venom is a potentially lethal and complex mixture of hundreds of functional toxins that vary extensively among snake species (5, 6). Although snake venom exhibits major variations, the various toxins present can be broadly classified into causing three major categories of pathology in snakebite victims: haemotoxicity, neurotoxicity and cytotoxicity (7).

Envenomings by viperid snakes (Viperinae, true vipers; Crotalinae, pit vipers) are predominately responsible for causing haemotoxicity, and are systemically characterised by overt haemorrhage, such as from the gums, and internal haemorrhage, such as intracranially (7, 8). Snakebite induced coagulopathy, i.e., defibrinogenation, also known as venom-induced consumption coagulopathy (VICC), is one of the most common life-threatening and important systemic pathologies observed following snakebite (2, 9-11). VICC is caused by the bites of a wide range of snakes, including many true vipers and pit vipers (e.g., *Daboia russelii, Callosalesma rhodostoma*, etc), certain Australasian elapids (e.g., *Oxyuranus scutellatus*) and a few colubrid snakes (e.g., *Dispholidus typus*) (12, 13).

The venom toxins primarily thought to be responsible for causing VICC are members of the snake venom metalloproteinase (SVMP) and snake venom serine protease (SVSP) toxin families (14, 15). Each of these toxin families is multi-locus in nature, in that related genes produce multiple related protein isoforms in the venom glands of snake species and, due to isoform diversity as the result of protein neofunctionalisation, these isoforms typically exhibit distinct functional activities (7, 13, 16). Snakebite coagulopathy disorders result from the consumption of clotting factors (e.g., Factor X, V and prothrombin) (15) via initial toxin-induced cleavage stimulating activation of the coagulation cascade, resulting in the abnormal and continual activation of downstream clotting factors, and a loss of clotting capability via depletion of fibrinogen (13, 16). However, some toxins also act directly on fibrinogen in a fibrinogenolytic manner (15), and many venoms contain both upstream clotting factor activators and fibrinogenolytic toxins simultaneously. Thus, venom toxins can result in hypofibrinogenaemia, with either no formation of fibrin due to the cleavage of the fibrinogen α-chain and/or β-chain by thrombin-like enzymes or fibrinogenases (typically SVSP toxins), or afibrinogenaemia (the absence of circulating fibrinogen) due to depletion via upstream coagulation cascade activation (17). While this combination of procoagulant and anticoagulant toxins present in snake venom (18) can rapidly lead to a loss of clotting capability in envenomed victims (9, 15, 17), the severity of envenoming can be further exacerbated by other toxins (e.g., SVMPs) simultaneously stimulating widespread haemorrhage by disrupting the integrity of the microvasculature via cleavage of extracellular matrix proteins (19, 20).

Antivenoms, which consist of animal-derived polyclonal antibodies derived from hyperimmunised serum/plasma, remain the only specific therapeutic to combat the toxicity of snakebite envenoming (21). Although antivenoms are life-saving therapeutics, they are also associated with several limitations that affect their efficacy, safety, and utility in the tropical world. For example, the intravenous delivery of animal-derived antibodies comes with an associated risk of adverse reactions, which may include vomiting, urticaria, generalised rash or, in rarer cases, more severe effects (e.g., anaphylactic shock) (22-25). In addition, only 10-20% of antivenom antibodies are typically specific to the venom immunogens (26), as venom-immunised animals are exposed to other environmental stimuli and toxin-specificity is typically not selected for from the resulting antibody pool. Perhaps most importantly, antivenoms exhibit limited cross-snake species efficacy as the direct result of variation in venom composition among medically important snake species (6, 27). Thus, antivenom efficacy is largely restricted to the snake species, or those closely-related to or with similar venom compositions to, those used during the immunisation process (28). The result of limited paraspecific antivenom efficacy is a fragmented drug market, with numerous antivenoms manufactured around the world with specificity against certain snake species and thus restricted for use in specific geographical regions. Nonetheless, antivenoms are often unaffordable to patients, as treatment courses frequently necessitate the administration of multiple vials (ranging from 3-30 vials depending on region and product), resulting in costs exceeding USD 1,000 in parts of sub-Saharan Africa, for example (29). This cost pushes many tropical snakebite victims further into poverty (30, 31).

In response to the above, much recent research has focused on applying alternate therapeutic strategies to circumvent the limitations currently associated with conventional antivenoms. Promising approaches explored to date include the use of toxin-specific monoclonal antibodies (32), small-molecule drugs (33), decoy receptor binding proteins (34) and aptamers (35-37). Although arguably the least studied of these approaches in the context of snakebite, single-stranded DNA (ssDNA) aptamers have emerged as promising alternatives to antibodies in various biosensing platforms and as the biological recognition element for therapeutic applications and diagnostic tools for different diseases (38-40). Aptamers are short ssDNA (or ssRNA) molecules that can bind to their target and fold into complex and stable three-dimensional shapes (41, 42) with high specificity and affinity through hydrogen bonding, van der Waals forces, hydrophobic, salt bridges and other electrostatic interactions (43-46). Consequently, aptamers act like antibodies by binding and inhibiting target antigens, and thus have been referred to as chemical antibodies due to their synthetic production (47, 48). ssDNA aptamers possess a number of advantages over antibodies as, in addition to seemingly exhibiting comparable high-affinity recognition properties (i.e., dissociation constants in the low nanomolar (nM)/picomolar (pM) range) (43, 44, 49), they are highly specific, sensitive, chemically stable with long shelf lives (50-52), and require facile artificial synthesis (53) resulting in high purity, low inter-batch variability and cost-effective production (54-56). Further, ssDNA aptamers are poorly immunogenic which aids their tolerance, and due to their small size they diffuse readily into tissues (57). However, this latter characteristic also causes challenges, as the *in vivo* half-life of ssDNA aptamers is much shorter (∼20 minutes) than antibodies (e.g., potentially a few days to weeks) (58), though prolonged circulation can be improved by chemical modification (57, 59-61). Despite the therapeutic promise of aptamers, to date the only FDA approved aptamer remains the macular degeneration treatment Macugen® (Pegaptanib) which was first licenced in 2005 (62). Since then, no other aptamers have been entered the market, although many candidate aptamers have entered clinical trials over the past decade, including as promising new treatments for heart disease, cancer and type II diabetes (62).

ssDNA aptamers are typically selected by Systematic Evolution of Ligands by Exponential enrichment (SELEX) technology, which was introduced by Ellington and Tuerk three decades ago (43, 44). This process is based on the isolation of high-affinity ligands from a combinatorial ssDNA library (approximately 10^17^ random sequence oligonucleotides) through repeated cycles (usually requiring seven to fifteen rounds) of binding, elution, and amplification to select for oligonucleotides with desired specificity and/or sensitivity (48, 63, 64). Following cyclical SELEX enrichment, the final ssDNA aptamer pool is subjected to sequencing to identify the optimal resulting binding sequences, which are then manufactured by chemical synthesis and downstream optimised by introduction of chemical modifications to further improve their pharmacokinetic and/or pharmacodynamic profiles (65). To date, several ssDNA aptamers have been successfully selected for and applied against a wide range of targets that include, proteins, peptides, small organic compounds, carbohydrates, metal ions and biological cofactors (66-72). In addition, ssDNA aptamers have been shown to bind to highly toxic antigens, which, due to toxicity, cannot be achieved in animal-based methods to generate specific antibodies (73). In the context of animal toxins, little work has been done, though aptamers have been selected against the three-finger toxin α-bungarotoxin from *Bungarus multicinctus* (74) and shown to exhibit cross-recognition and inhibitory activity against cytotoxins from *Naja atra* venom that share similar tertiary structures (37). Another recent study selected ssDNA aptamers against Indian *Bungarus caeruleus* (common krait) venom and used them in a diagnostic context to discriminate envenomings by this species from those of other medically important snakes found in the region (75). In this study, we sought to explore the potential utility of ssDNA aptamers against snake venom SVSP toxins responsible for contributing to venom induced coagulopathy following snakebite envenoming. Using SELEX technology, we selected ssDNA aptamers against recombinantly expressed versions of the fibrinogenolytic SVSPs Ancrod from the venom of *Calloselasma rhodostoma* and Batroxobin from *Bothrops atrox*. Downstream ssDNA Aptamer Linked Immobilised Sorbent Assay (ALISA), fibrinogenolysis, and coagulation experiments demonstrated that the rationally selected ssDNA aptamers were able to recognise native and recombinant SVSP toxins and inhibit toxin and venom-induced prolongation of plasma clotting times and consumption of fibrinogen. These data highlight the potential utility of ssDNA aptamers as novel toxin-inhibiting therapeutics and alternative binding scaffolds for future treatment strategies targeting tropical snakebites.

## Materials and methods

### ssDNA library and primer design

The ssDNA library and PCR primers were chemically synthesised and purified by Integrated DNA Technologies, Inc. (90). The random ssDNA library used in the first SELEX cycle consisted of (3 nM of 2 × 10^17^ sequences) nucleotides. The ssDNA library consisted of a central random region of 72 nucleotides flanked by two primers consisting of 16 nucleotides at the 5’ and 3’ end (5’-TCCCTACGGCGCTAAC-N72-GTTGTAGCACGGTGGC-3’) for amplification of the library sequence. To facilitate the separation of ssDNA from amplified double-stranded (dsDNA) PCR products and to quantify DNA during selection, the forward and reverse primers were designed and modified with fluorescein and a HEG linker (the spacer is designed to block the PCR extension step during amplification). The modified PCR forward primer used during the selection cycles was (5’-6-FAM-TCCCTACGGCGCTAAC-3’), and the reverse primer was (5’-poly dA20-HEG-spacer-GTTGTAGCACGGTGGC-3’). An unmodified primer set was used for PCR amplification and cloning when SELEX rounds were completed and enriched.

### Target conjugation

The recombinant toxins used in this study, Ancrod from *C. rhodostoma* and Batroxobin from *B. atrox*, were previously expressed in HEK293 mammalian cell lines (76). In this study, we used these toxins to conjugate to NHS-activated Sepharose® 4 Fast Flow beads (Sigma-Aldrich, UK) to generate ssDNA aptamers as novel toxin inhibiting therapeutics. To do so, the manufacturer protocol was followed, with first a 300 μg/ml solution of each toxin prepared in coupling buffer (0.1 M NaHCO_3_, 0.5 M NaCl, pH 8.3). Next, 2 mL of NHS-activated Sepharose beads were rinsed multiple times with 1 mM HCl for 15 minutes to remove the additives and preserve the activity of the reactive groups. Each venom toxin was then added immediately to the washed Sepharose beads independently (1:1 v:v ratio) and mixed by end-over-end rotation overnight at 4 °C. After incubation, the beads were centrifuged at 14 x g for 5 minutes, the supernatant discarded, and the samples washed with 1 ml of 1 M NaCl. This centrifugation step was repeated before 2 ml of 1 M NaCl was added, and the samples were incubated with rotation for 2 hours at 4 °C to block the unreacted amine groups on the beads. After blocking, the toxin-conjugated Sepharose was washed three times with 4 ml in an alternating manner, first with 0.1 M acetic acid/sodium acetate, pH 4 containing 0.5 M NaCl 0.1 M, followed by Tris-HCl buffer, pH 8 containing 0.5 M NaCl. Thereafter, the toxin-conjugated Sepharose was stored in 50 mM Tris-HCl buffer (pH 7.5), 0.05% sodium azide at 4 °C until use.

### PCR amplification

Each of the recombinant toxin-conjugated Sepharose mixtures were washed five times by adding 500 μl binding buffer (50 mM Tris, pH 7.5, 150 mM NaCl, 2 mM MgCl_2_) to 100 μl of Sepharose, using centrifugal tube filters (0.45 µm) (Costar, USA). In the first SELEX cycle, 50 μl (3 nM) of the ssDNA library solution in 450 μl binding buffer was used, whereas 50 pM of the DNA elution from each round was used for subsequent cycles. The DNA and binding buffer mixture were heated to 90 °C on a heat block for five minutes, cooled at 4 °C for ten minutes and then held at room temperature (RT) for five minutes. Next, 300 μl of DNA and binding buffer mixture was added to the washed beads and incubated end-over-end for 1 hour at RT. After incubation, the mixtures were washed and centrifuged five times with 500 μl of binding buffer at 82 x g for 2 minutes, to ensure the elimination of unbound DNA. The flow-through from the first wash was retained for use as a positive control in downstream agarose and denaturing polyacrylamide gel electrophoresis (PAGE) experiments. DNA bound to the toxin-conjugated beads was collected from each sample using elution buffer (7 M urea in binding buffer) and incubated at 90 °C for 10 minutes on a heat block. The elution step has performed a total of five times via the addition of 300 μl elution buffer until no fluorescence was detected in the eluted aliquots, indicating the complete elution of bound DNA from the beads. The eluted aliquots were then desalted with filtered water and concentrated using an ultrafiltration device with a 3 kDa cut-off membrane (Amicon Ultra-0.5 Centrifugal Filter Unit; Merck, UK) to remove any urea and salt residues.

A double-stranded (dsDNA) product with different lengths for each strand was generated via PCR amplification by using two parallel 50 μl reactions (one for each toxin target), with each containing five units of Taq Plus and polymerase buffer (New England, BioLabs®, UK), 2 mM MgCl_2_, 200 μM dNTPs and 0.2 μM of the unmodified forward and reverse primers. The PCR conditions were: 94 °C for 10 minutes, followed by 17 cycles of 94 °C for 1 minute, 54 °C for 1 minute and 72 °C for 1 minute, and a final extension step of 10 minutes at 72 °C. To confirm the amplification of the dsDNA product, the samples were then analysed electrophoretically using 2% agarose gel electrophoresis.

A ssDNA product with different lengths for each strand was also generated via PCR amplification by using 20 parallel 50 μL reactions, using the same conditions as outlined above, except with the use of 0.2 μM of the labelled forward primer. Following amplification, the 20 reactions were pooled and aliquoted into three Eppendorf tubes, each containing 250 μl of pooled ssDNA product, 50 μl 3M sodium acetate, pH 5.2 and 750 μl absolute ethanol. The tubes were then mixed and stored at –80 °C for 2 hours, before centrifugation at 13,000 x g for 30 minutes at 4 °C. The supernatant was discarded, and the remaining pellet was dried at 90 °C for a couple of minutes on a heat block before resuspension in 400 μl of water and formamide (1:1 v/v) and incubation at 90 °C for 5 minutes.

### Separation of ssDNA by denaturing PAGE

We employed a 10% denaturing PAGE to separate the resulting ssDNA from the double-stranded (dsDNA) amplified PCR products. Fifty microlitres of each dsDNA sample was independently loaded into the gel and run at 200 V for 55 minutes using a Mini-PROTEAN Electrophoresis System (Bio-Rad, UK). Next, the ssDNA band was observed under UV light, dissected, cut and transferred to a 15 ml falcon tube containing 4 ml of Tris-Ethylenediaminetetraacetic acid (EDTA) buffer solution (TE buffer; 10 mM Tris, pH 7.4, 1 mM EDTA). Samples were then subjected to freeze-thaw for 30 minutes at −80 °C, 10 minutes at 90 °C, followed by incubation on a rotary shaker overnight at 37 °C (77). Finally, ssDNA was eluted and concentrated using an ultrafiltration device with a 3 kDa cut-off membrane (Amicon Ultra-0.5 Centrifugal Filter Unit; Merck, UK), as described earlier. The UV and the fluorescence intensity of each dsDNA and ssDNA were then measured for each eluted fraction by using a NanoDrop 2000C spectrophotometer and a FLUOstar Omega 384 well black assay plate reader (B.M.G. LabTech), respectively. The fluorescence spectra were detected at an excitation wavelength of 470 nm and an emission of 515 nm.

### Counter-selection

Following enrichment, counter (negative) selection was employed by using blank Sepharose beads instead of those conjugated to toxins. The goal here was to increase enrichment to the toxin target via elimination of aptamers directed towards the beads. Sepharose beads were subjected to the aforementioned conjugation process with 0.1 M Tris-HCl buffer, pH 8.3 containing 0.5 M NaCl only (i.e., in the absence of toxins), and counter selection followed the same experimental strategy as described above and was used to filter out negative targets.

### Cloning and sequencing ssDNA aptamers

After ssDNA recovery, the eluted ssDNA from the 14^th^ round of selection was amplified using the unlabelled PCR primer (5’-TCCCTACGGCGCTAAC-3’) following the same conditions as PCR amplification: 94 °C for 10 minutes, followed by 17 cycles of 94 °C for 1 minute, 54 °C for 1 minute and 72 °C for 1 minute, and a final extension step of 10 minutes at 72 °C. Next, ligation was performed by using the TOPO TA Cloning Kit for Sequencing and One Shot TOP10 Chemically Competent *E. coli* (Thermo-Fisher Scientific, UK). Ligation reactions consisted of 2 µL of fresh PCR product, 1 µL of salt solution, 1 µL of TOPO® vector and 1 µL of nuclease-free water, with the resulting samples mixed gently and incubated for 30 minutes at RT (20-25 °C). For cloning, 5 µL of the ligated product was added to a vial of One Shot TOP10 Chemically Competent *E. coli* (Thermo-Fisher Scientific, UK) and mixed gently. The cells were then placed on ice for 30 minutes, heat shock treated for 30 seconds at 42 °C and transferred back to ice. Next, 250 µL of pre-warmed S.O.C medium was added, and samples were incubated at 37 °C for 1 hour with horizontal shaking at 200 rpm. Thereafter, 50 µL of the medium was spread on 25 ml of Lauria Broth (LB) agar (VWR, UK) plates supplemented with 25 µL ampicillin (20 mg/ml) (Sigma-Aldrich, UK), 250 µL X-Gal (5-bromo-4-chloro-3-indolyl-β-D-galactoside) (20 mg/ml) (Sigma-Aldrich, UK) and Isopropyl ß-D-1-thiogalactopyranoside (IPTG) 25 µL (100 mM) (Sigma-Aldrich, UK), and incubated at 37 °C overnight. Following incubation, ten random positive colonies (white colonies) were picked for colony PCR analysis: 5 µL 10x PCR buffer, 0.5 µL deoxyribonucleoside triphosphates (dNTPs) mix (50 mM), 0.5 µL of forward and reverse M13 primers (0.1 µg/µL each), 41.5 µL nuclease-free water and 1 µL Taq polymerase (1 unit/µL). Colony PCR was performed using the following cycling parameters: initial denaturation at 94 °C for 2 minutes, followed by 25 cycles of denaturation at 94 °C for 1 minute, annealing at 55 °C for 1 minute and extension at 72 °C for 1 minute, with a final extension step at 72 °C for 7 minutes. To confirm insert amplification, 2% agarose gel electrophoresis was performed.

One hundred positive bacterial colonies were sterilely selected and inoculated with 5 ml of LB medium containing 50 µL Ampicillin (20 mg/ml) and incubated overnight at 37 °C with shaking at 200 rpm. Next, plasmid DNA was purified using a QIAprep Spin Miniprep Kit (Qiagen, Germany). The overnight culture was centrifuged at 6000 x g for 10 minutes at RT. The pellet was then resuspended in 250 µl Buffer P1 (50 mM Tris-HCl pH 8.0, 10 mM EDTA, 100 μg/ml RnaseA) and transferred to a microcentrifuge tube. Then, 250 µl Buffer P2 (lysis buffer: 200 mM NaOH, 1%) was added and mixed thoroughly by inverting the tube gently 4–6 times. After that, 350 μl Buffer N3 was added and mixed immediately and thoroughly by inverting the tube 4–6 times. Next, samples were centrifuged for 10 minutes at 17,900 x g, and the supernatant was transferred to QIAprep 2.0 Spin Columns before brief centrifugation at 2,817 x g for one minute with the eluate discarded. Columns were then washed with 750 µl Buffer PE via centrifugation at 2,817 x g for one minute, the eluate discarded, and columns centrifuged for an additional minute to remove any residual buffer. Finally, 50 μl Buffer EB (10 mM Tris-HCl, pH 8.5) was added to each column and incubated at RT for two minutes prior to elution via centrifugation at 2,817 x g for 1 minute. The resulting PCR products were then sequenced commercially via Sanger sequencing (SourceBioScience, UK), and the resulting ssDNA aptamer sequences were analysed and aligned using the PRALINE website: (https://www.ibi.vu.nl/programs/pralinewww/).

### Dissociation constants (K_dS_)

The binding affinities of the identified ssDNA aptamers against each of the corresponding toxins used for selection were evaluated using a fluorescence-based assay. Both forward and reverse primer-binding sites were truncated and labelled with 6-FAM (fluorescein-labelled) before synthesis. Different concentrations of fluorescein-labelled ssDNA aptamer (0, 5, 10, 50, 100, 200, 300 and 400 nM) were incubated with a constant amount of the corresponding toxin-conjugated beads (50 µL) as described in the selection protocol (by heating to 90 °C for 10 minutes, 4 °C for 10 minutes and then at RT for 5 minutes. After 1 hour of incubation at RT with rotation, the unbound ssDNA aptamers were washed using 3 kDa centrifuge filters and 500 µL binding buffer, and the toxin-bound ssDNA aptamers were eluted five times via the addition of 300 μl elution buffer (7 M urea in binding buffer) and incubation at 90 °C for 10 minutes on a heat block. The five eluted aliquots were then concentrated using an ultrafiltration device with a 3 kDa cut-off membrane (Amicon Ultra-0.5 Centrifugal Filter Unit; Merck, UK) and desalted with filtered water to remove urea and salts. The fluorescence intensity of each ssDNA was then measured for each eluted fraction at an excitation wavelength of 470 nm and emission of 515 nm. The K_d_ of each ssDNA aptamer was calculated via nonlinear regression analysis of the resulting plotted hyperbolic curves using Prism v8 software (GraphPad).

### Binding by ALISA

To quantify reciprocal binding to the recombinant toxins (Ancrod and Batroxobin) by the highest affinity aptamers (determined by the K_d_ analysis above) we used an ALISA assay. We also assessed binding to native venom samples that corresponded to the source of each toxin (i.e., *Calloselasma rhodostoma*, captive bred and *Bothrops atrox*, Brazil) and an additional panel of eleven geographically and taxonomically diversity snake species (*Bitis arietans*, Nigeria; *Bothrops asper*, Costa Rica; *Bothrops jararaca*, Brazil; *Crotalus atrox*, captive bred; *Daboia russelii*, Sri Lanka; *Deinakgistrodon acutus*, captive bred; *Dispholidus typus*, South Africa; *Echis carinatus*, India; *Echis ocellatus*, Nigeria; *Rhabdophis subminiatus*, Hong Kong; *Trimeresurus albolabris*, captive bred) to assess breadth of binding. Many of these venoms were collected from animals maintained under controlled environmental and dietary conditions in the UK Home Office licensed and inspected herpetarium of the Centre for Snakebite Research and Interventions (CSRI) at the Liverpool School of Tropical Medicine. The remainder were sourced from historical venom collections held in the CSRI herpetarium (specifically, *Bothrops asper, Bothrops jararaca*, Calloselasma *rhodostoma, Daboia russelii, Rhabdophis subminiatus* and *Trimeresurus albolabris*) or commercially purchased from Latoxan, France (*Dispholidus typus*). Lyophilised venoms were stored at 4 °C and reconstituted with phosphate-buffered saline (PBS) (0.12 M NaCl, 0.04 M phosphate, pH 7.2) to a concentration of 1 mg/ml prior to use.

For the ALISA, 1 µl of Modifier reagent (HRP Conjugation Kit - Lightning-Link®, ab102890) was added to 10 µl of each ssDNA aptamer (Ancrod-55 and Batroxobin-26; 200 nM), mixed gently and left standing for 3 hours in the dark at RT (20-25 °C). After incubation, 1 µl of Quencher reagent (HRP Conjugation Kit -Lightning-Link®, ab102890) was added to each ssDNA aptamer sample and mixed gently. Next, microtiter 96 well ELISA plates (Thermo Fisher Scientific) were coated with coating buffer (100 mM carbonate/Bicarbonate buffer, pH 9.6) containing 100 ng of each toxin or native venom and incubated overnight at 4 °C. Plates were then washed three times with PBST (1 x PBS, pH 7.4 containing 0.1% Tween), before blocking with 300 µl/well of 0.5% bovine serum albumin (BSA; Sigma-Aldrich, UK) diluted in PBS (filtered with MF-Millipore® Membrane Filter, 0.45 µm pore size). Following incubation at RT for two hours, the plates were washed another three times with PBST. Next, two concentrations of the aptamers were prepared and diluted in nuclease free water (2 nM and 0.4 nM). Then, 100 µl of each aptamer, alongside water (as a negative control), were added to the plate in duplicate, followed by incubation for 2 hours at RT. The plates were then rewashed three times with PBST before the addition of 3,3’,5,5’-Tetramethylbenzidine (TMB) Liquid Substrate System for ELISA (Sigma-Aldrich, UK) and incubated for 12 minutes in the dark at RT. To stop the reaction, 25 µl of 20 % Sulfuric acid was added, before the signal was read spectrophotometrically at 450 nm on a FLUOstar Omega microplate reader. Finally, all reads at 450 nm were subtracted from 540 nm to remove any background signal.

### Fibrinogenolysis via degradation SDS-PAGE

To assess the capability of the selected ssDNA aptamers to inhibit the fibrinogenolytic activity of the recombinant toxins and corresponding native snake venoms, we used a degradation SDS-PAGE (sodium dodecyl sulfate polyacrylamide gel electrophoresis) approach. The toxins Ancrod and Batroxobin (7 μg, 1 mg/ml), and the corresponding venoms from *C. rhodostoma* and *B. atrox* (7 μg, 0.3 mg/ml), were incubated with human plasma fibrinogen (3 µg, 2.5 mg/ml, Sigma-Aldrich, UK) for two hours at 37 °C. All immunogen samples were also used in neutralisation experiments following a pre-incubation step at 37 °C for 15 minutes with each of the selected ssDNA aptamers (1 pM). Next, all samples were incubated for two hours at 37 °C. Human plasma fibrinogen with PBS instead of venom was used as the negative control. After incubation, 10 µl of each sample was mixed at a 1:1 volume/volume with reducing buffer (3.55 µl H_2_O, 1.25 ml 0.5 M Tris (pH 6.8), 2.50 ml glycerol, 2.0 ml 10% SDS, 1.50 ml saturated bromophenol blue and 150 µl β-mercaptoethanol) and heated for 5 minutes at 100 °C. For gel electrophoresis, ten well 8% SDS-PAGE gels were hand-cast, using the following approach: 10 ml resolving gel (4.7 ml H_2_O, 2.5 ml Tris pH 8.8 (1.5 M), 2.7 ml 30% bis-acrylamide, 50 µl 20% SDS, 100 µl 10% ammonium persulfate (APS) and 7 µl tetramethylethylenediamine (TEMED)); 4 ml of 4% stacking gel (2.7 ml H_2_O, 0.5 ml Tris pH 6.8 (1 M), 800 µl 30% bis-acrylamide, 20 µl 20% SDS, 40 µl 10% APS and 4 µl TEMED). Next, 10 µl of each sample, alongside 5 µl of broad molecular weight protein marker (Broad Range Molecular Marker, Promega), was loaded onto the gel and run at 200 volts for 55 minutes using a Mini-PROTEAN Electrophoresis System (Bio-Rad, UK). Resulting gels were then stained at a final concentration of 0.1% (w/v) Coomassie blue R350 (0.4g of Coomassie blue R350 in 200 mL of 4% (v/v) methanol in H_2_O), 10% (v/v) acetic acid and 20% (v/v) methanol overnight at RT. Gels were destained (4:1:5 methanol: glacial acetic acid:H_2_O) for at least two hours at RT. For visualisation, gels were subsequently imaged using a Gel Doc EZ Gel Documentation System (Bio-Rad, UK).

### Sample preparation for clotting profiling experiments

Blood samples for clotting profiling experiments were obtained according to ethically approved protocols (LSTM research tissue bank, REC ref. 11/H1002/9) from consenting healthy volunteers who confirmed they had not taken any anticoagulant treatments for at least three months prior to blood collection. Blood samples were collected in tubes containing acid citrate dextrose adenine (ACD-A) as an anticoagulant. To prepare Fresh Frozen Plasma (FFP) a fresh blood sample was centrifuged at 2500 × g at 20-25 °C for 10 minutes, and the supernatant was retained and stored at −80 °C until use. For Platelet Poor Plasma (PPP), FFP samples were centrifuged again in an identical manner, with the resulting platelet-depleted supernatant stored at −80 °C until further use. To prepare cryo-precipitate Anti Haemolytic Factor (Cryo-AHF), FFP stored at −80 °C (for at least 24 hours) was slowly thawed at 2-6 °C for 30-45 minutes. Precipitated proteins were immediately sedimented in a refrigerated centrifuge (2-6 °C) at 5000 × g for 20 minutes. Next, the supernatant, except a ∼500 µl layer, was carefully removed, and the retained supernatant layer was then mixed gently with the precipitate, and the resulting Cryo-AHF samples were either stored at −20 °C until use or, in the case of immediate use, 1 ml of Cryo-AHF was suspended in 2 ml of 0.9 % saline solution (NaCl) with short terms storage (<48 hours) at 4 °C (78).

In addition to the samples prepared above, we also used commercially sourced samples (Diagnostic Reagents Ltd, UK) to serve as quality control checks for the various reagents employed. To this end, normal and abnormal test controls were implemented for the experiments described below prior to experimentation. These consisted of 1) Diagen control plasma for fibrinogen: normal (RCPN070) and abnormal (RCPA080) and 2) control plasma for aPTT and PT: normal (IQCN130), abnormal 1 (Mild, IQCM140) and abnormal 2 (Severe, IQCS150). All QC samples were aliquoted into 500 µl, stored at - 80 °C, and defrosted in a water bath for 5 minutes at 37 °C immediately prior to use. To act as positive controls for the aptamers, we used commercial antivenoms, specifically the Thai Red Cross monovalent equine Malayan Pit Viper antivenom (Lot #: CR00316, expiry date 06/2021) for Ancrod and *C. rhodostoma* experiments and the Instituto Butantan polyvalent equine SORO antibotropico/crotalico antivenom (Lot #: 1012308, expiry date: 2013) for Batroxobin and *B. atrox* experiments. These freeze-dried antivenoms were reconstituted in pharmaceutical grade water supplied by the manufacturers, the protein concentrations quantified using a Nanodrop (Thermo-Fisher Scientific, UK), and then stored at 4 °C short terms until use.

### Fibrinogen consumption via the Clauss method

The Clauss method is a quantitative, clot-based assay. It measures the ability of thrombin to convert fibrinogen to fibrin clot followed by manual time measurements of clotting (79). Here we applied this method in toxin- and venom-spiking experiments to assess the inhibitory capability of the selected ssDNA aptamers against the depletion of fibrinogen. Twenty microlitres of either FFP, PPP or AHF were spiked with 0.6 ng of Ancrod, Batroxobin, *C. rhodostoma* venom or *B. atrox* venom, or 0.9 % saline solution as the negative control. All samples were also repeated in the presence of 1 pM of the selected ssDNA aptamers or 0.5 µg of the commercial antivenoms following preincubation at 37 °C for 5 minutes. Samples were then diluted tenfold with 0.02 M imidazole buffer (pH 7.35), transferred to glass test tubes (10 × 75 mm), and warmed at 37 °C for 120 seconds. Thereafter, 100 µl of thrombin reagent (20 units/ml) (Diagnostic Reagents Ltd, UK) was added, and time measurements commenced. Tubes were gently tilted at regular intervals (returning to the water bath between tilting), and the time for the formation of a clot recorded. All experiments were performed in duplicate.

### Prothrombin Time (PT)

To measure the inhibitory capability of selected aptamers against toxins acting on the extrinsic coagulation pathway, we quantified differences in the PT between toxin and venom samples in the presence and absence of aptamers. Measurements of PT were undertaken by first adding 100 µl of Calcium Rabbit Brain Thromboplastin (Diagnostic Reagents Ltd, UK) to a glass test tube (10 × 75 mm) and incubating at 37 °C for 60-120 seconds in a water bath. Next, 50 µl of PPP or cryo-AHF was spiked with 50 µl containing 0.6 ng of Ancrod, Batroxobin, *C. rhodostoma* venom or *B. atrox* venom, or saline solution as the negative control. All samples were also repeated in the presence of 1 pM of the selected ssDNA aptamers or 0.5 µg of the commercial antivenoms, following preincubation at 37 °C for 5 minutes. Time measurements were commenced, tubes were gently tilted at regular intervals (returning to the water bath between tilting), and the time for the formation of a clot was recorded. All experiments were performed in duplicate. We did not use FFP for quantification of the PT as abundant platelets have been shown to interact with test reagents and increase the concentration of phospholipids, resulting in false shortening of clotting times (80).

### Activated Partial Thromboplastin Time (aPTT)

To measure the inhibitory capability of selected aptamers against toxins acting on the intrinsic and common coagulation pathways, we quantified differences in the aPTT between toxin and venom samples in the presence and absence of aptamers. To do so, 50 µL of Micronised Silica/Platelet Substitute Mixture (Diagnostic Reagents Ltd, UK) was placed in glass test tubes (10 × 75 mm) in a water bath at 37 °C and incubated for 60-120 seconds. Next, 50 µl of PPP or cryo-AHF was spiked with 0.6 ng of Ancrod, Batroxobin, *C. rhodostoma* venom or *B. atrox* venom, or saline solution as the negative control. All samples were also repeated in the presence of 1 pM of the selected ssDNA aptamers or 0.5 µg of the commercial antivenoms, following preincubation at 37 °C for 5 minutes. Thereafter, samples were added to the glass test tubes and gently tilted at regular intervals for precisely five minutes at 37 °C. Finally, 50 µL of 25 mM calcium chloride (pre-incubated at 37 °C) was added to each tube, and the tube was gently tilted until the resulting clot time was recorded. All experiments were performed in duplicate. As with PT, we did not use FFP for quantification of the aPTT due to the potential for interference by platelets.

## Results

### ssDNA aptamer selection

Here, we report the identification of ssDNA aptamers that target the recombinantly expressed SVSPs toxins Ancrod from *C. rhodostoma* and Batroxobin from *B. atrox*. To do so, we employed a parallel process consisting of 14 rounds of SELEX selections against each toxin target using a ssDNA library pool of 10^17^ random sequences. The selection process was monitored throughout by PCR amplification of the eluted DNA and agarose gel electrophoresis (Figure 1A and 1B), and by measuring the fluorescence intensity of the eluted ssDNA solution (Figure 1C and 1D).

**Figure 1.**
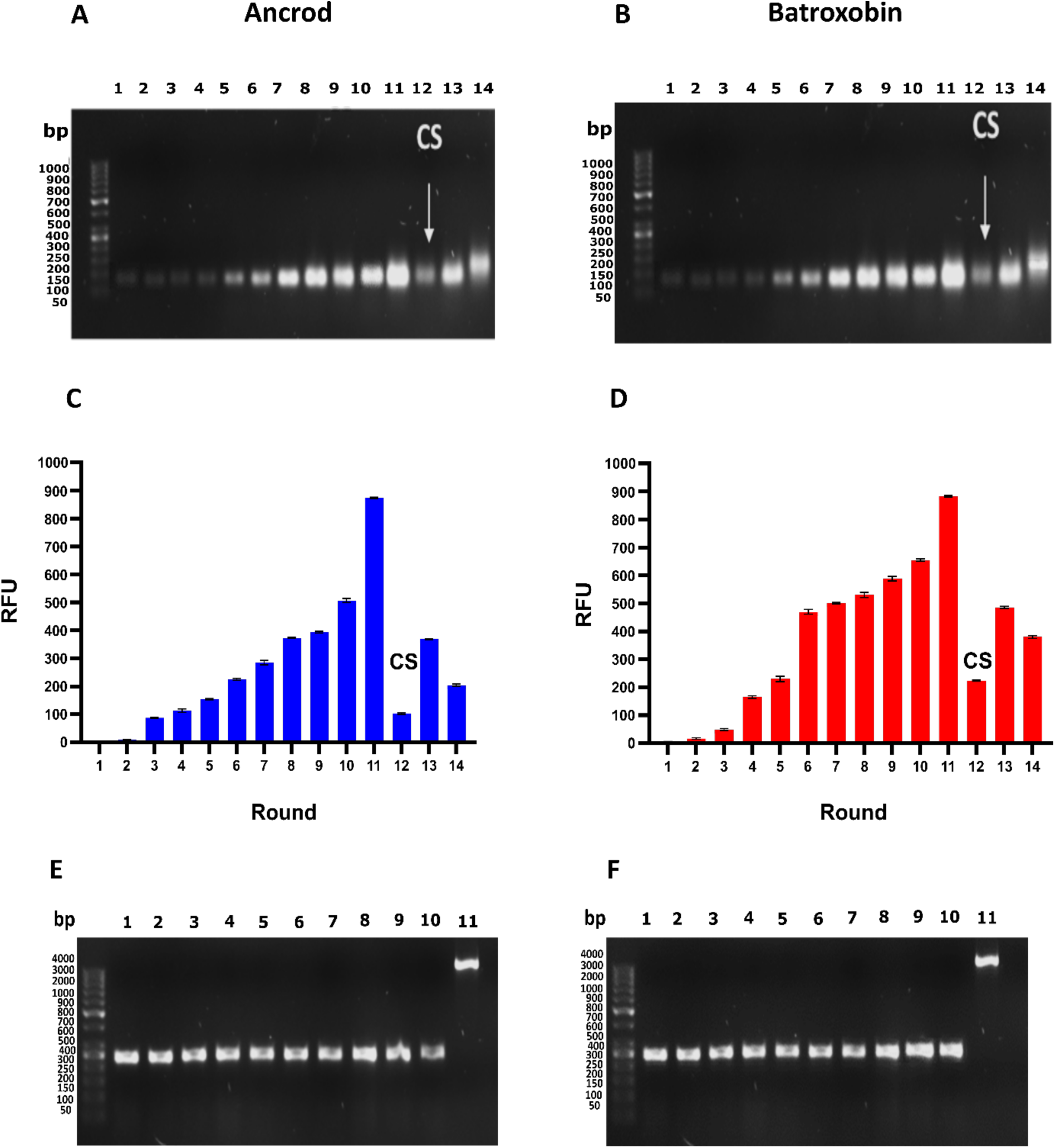
Sequential SELEX selection of aptamers against the recombinant SVSP toxins Ancrod and Batroxobin. **A** and **B)** DNA was amplified by PCR for each round of toxin selection employed (Lane 1-14; size, DNA≈104 bp), with non-specific bound ssDNA eliminated by counter selection (CS) employed at round 12. Data for Ancrod is shown in **(A)** and Batroxobin in **(B). C** and **D)** The fluorescence intensity of the eluted ssDNA solution from each round of selection was measured at excitation wavelength of 470 nm and emission of 515 nm. Error bars represent the standard error (SEM) of triplicate measurements, with data for Ancrod shown in **(C)** and Batroxobin in **(D). E** and **F)** Colony PCR was performed by ligation into a cloning vector and transformation into *E. coli* competent cells, with validation performed on ten random colonies to demonstrate that the inserted DNA (Lane 1-10; size≈300 bp) was successfully ligated into the pCR2.1-TOPO cloning vector (Lane 11; size≈3956 bp). Data for Ancrod is shown in **(E)** and Batroxobin in **(F)**.

Our findings revealed that initial DNA recovery against each target was low, though this increased with progressive rounds of selection cycles, until a marked increase was observed at round 11, indicating enrichment of the DNA pool with sequences specific to the toxins coupled to the beads (Figure 1). Notably, the patterns of fluorescent intensities obtained from the sequential selection rounds were highly comparable between the two toxin targets (Figure 1). Counter selection (CS) was performed at round 12 to eliminate non-specific ssDNA interacting with the sepharose beads instead of the toxins, and resulted in an anticipated drop in the fluorescence intensity of the recovered DNA (Figure 1C and 1D). Thereafter, two additional rounds of SELEX selection were performed to secondarily enrich toxin-specific aptamers, which resulted in an increase in fluorescence intensity at round 13, but no further increase at round 14, suggesting that target binding sites might be saturated (Figure 1C and 1D). Consequently, SELEX selection was stopped and the resulting ssDNA from the last round (round 14) was cloned for downstream aptamer identification.

Cloning was performed by ligation into a cloning vector and transformation into *E. coli* competent cells, with validation performed on ten random colonies to demonstrate that the inserted DNA (300 bp) was successfully ligated into the pCR2.1-TOPO cloning vector (3956 bp) (Figure 1E and 1F). Thereafter, 100 colonies were picked for each toxin target, and subjected to PCR followed by Sanger sequencing. Bioinformatic analyses of the resulting DNA identified various identical sequences, indicating enrichment of the DNA pool, and resulted in thirteen and nine unique aptamer sequences for Ancrod and Batroxobin, respectively (Table 1).

**Table 1.**
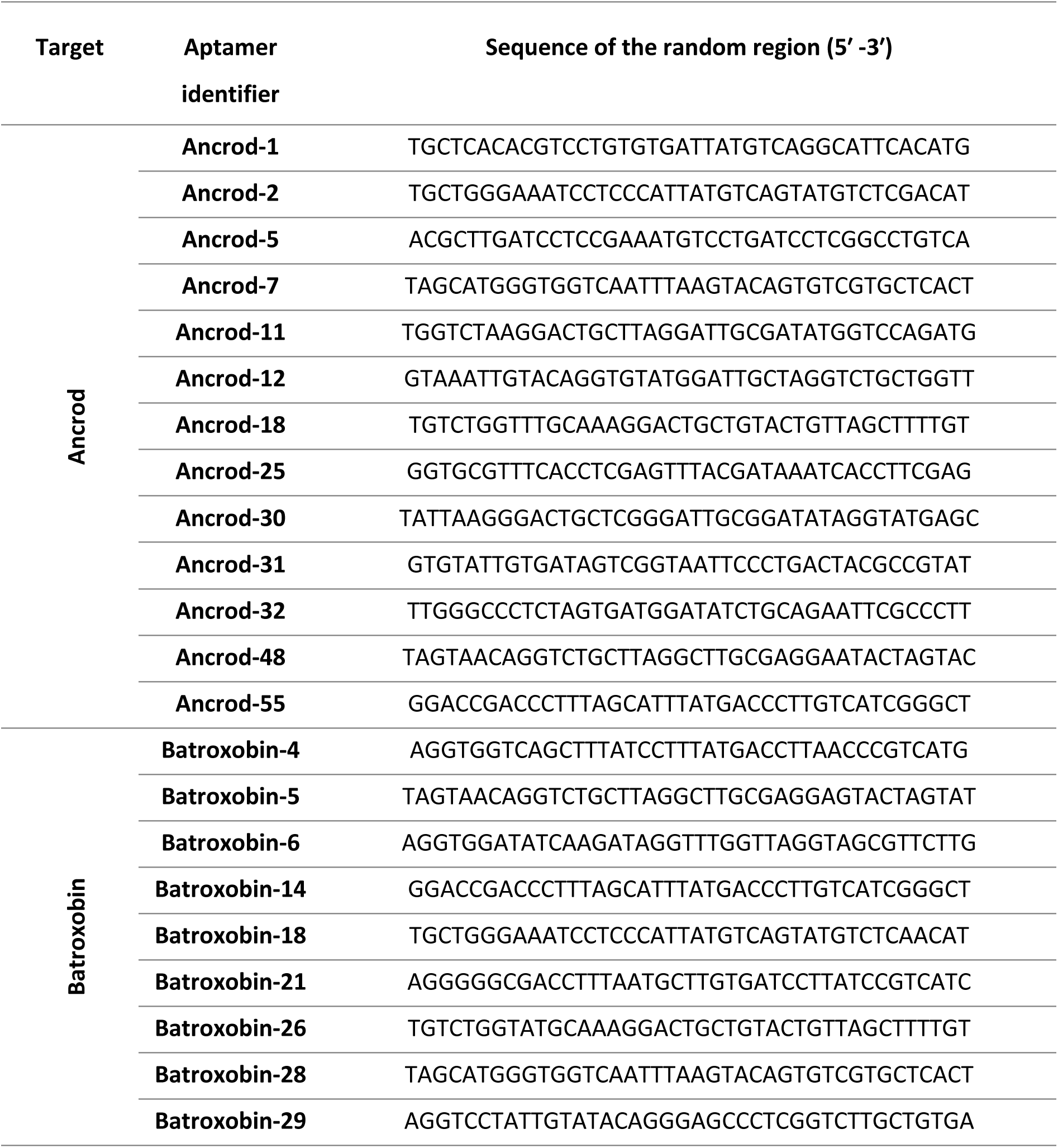
The identity of the unique ssDNA aptamers selected against Ancrod and Batroxobin.

### Dissociation constants (K_ds_) of identified aptamers

The dissociation constants (K_ds_) for each of the selected ssDNA aptamers, thirteen against Ancrod and nine against Batroxobin, were determined via nonlinear regression of saturation graphs resulting from concentration-based fluorescence binding assays. Of the 22 aptamers tested, only three ssDNA aptamer sequences directed against each of the targets exhibited nanomolar affinity, with those targeting Ancrod ranging from 3.0 nM to 17.8 nM and those against Batroxobin from 4.7 nM to 24.3 nM (Figure 2). The remaining ssDNA aptamers exhibited high K_d_ readings and were excluded from further study. The candidate ssDNA aptamer sequences with the lowest K_d_ were Ancrod-55 (K_d_= 3.0 nM) and Batroxobin-26 (K_d_= 4.7 nM) (Figure 2), and thus these two aptamers were selected for downstream recombinant toxin/crude venom binding and inhibition experiments.

**Figure 2.**
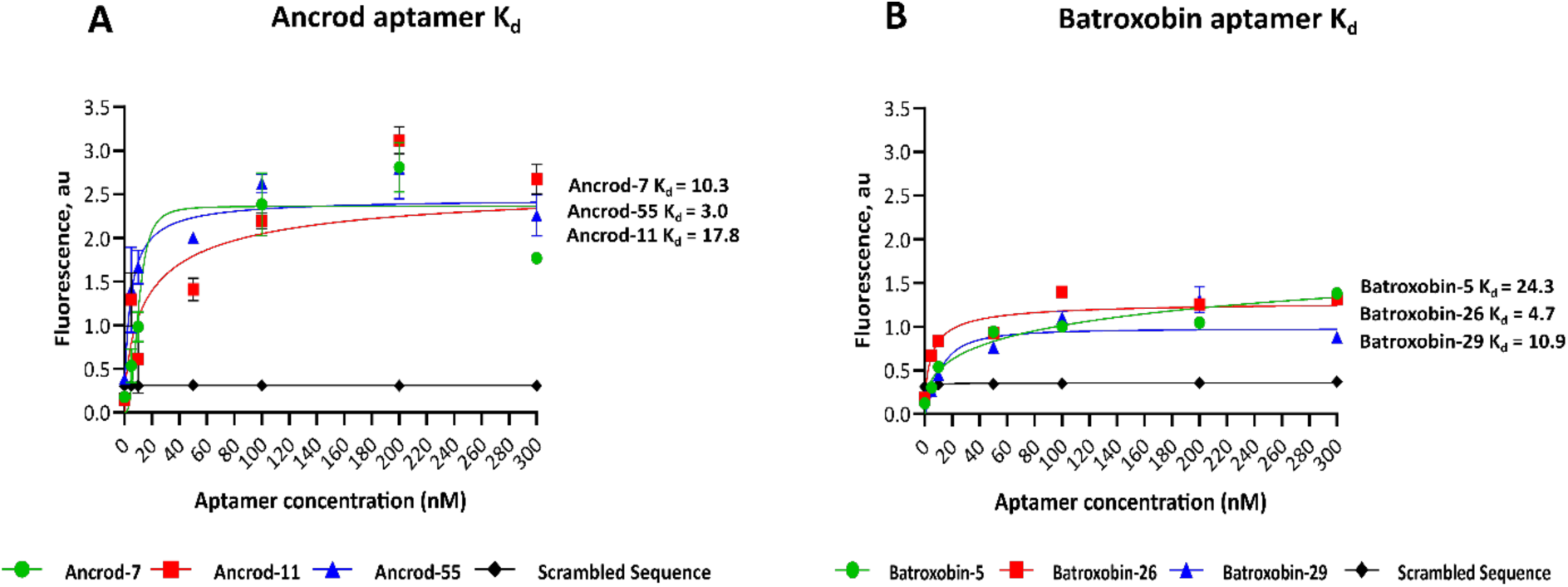
The equilibrium dissociation constants (K_d_) of the highest affinity ssDNA aptamers were derived from fluorescence binding curves for each toxin target. The data shown represents the highest affinity candidate aptamers against Ancrod (Ancrod-7, Ancrod-11 and Ancrod-55) **(A)** and Batroxobin (Batroxobin-5, Batroxobin-26 and Batroxobin-29) **(B)**. Concentration-dependent data was generated at an excitation wavelength of 470 nm and emission of 515 nm, and dissociation constants were calculated via nonlinear regression analysis of the resulting saturation curves. Data represent the mean of duplicate measurements, and error bars represent standard deviations (SD).

### Quantifying aptamer binding to recombinant toxins and venom by ALISA

An ALISA was next used to quantify the binding levels of the two highest affinity aptamers (Ancrod-55 and Batroxobin-26) against each of the recombinant toxins, and the native venoms from which the recombinant toxins were derived (i.e., *C. rhodostoma* and *B. atrox* venom) in a reciprocal and comparative manner. The results of these binding assays, which were performed at two aptamer concentrations (0.4 nM and 2 nM), revealed that the two aptamers tested exhibited considerable cross-recognition and binding to the two SVSP toxins (i.e., irrespective of which toxin the aptamer was selected against) and the two homologous venoms (Figure 3). Indeed, the aptamer Batroxobin-26 exhibited highly comparable binding levels to Batroxobin, Ancrod and the two native snake venoms (Figure 3B). These binding levels were, however, lower than those obtained with the Ancrod-55 aptamer for all comparisons (Figure 3A).

**Figure 3.**
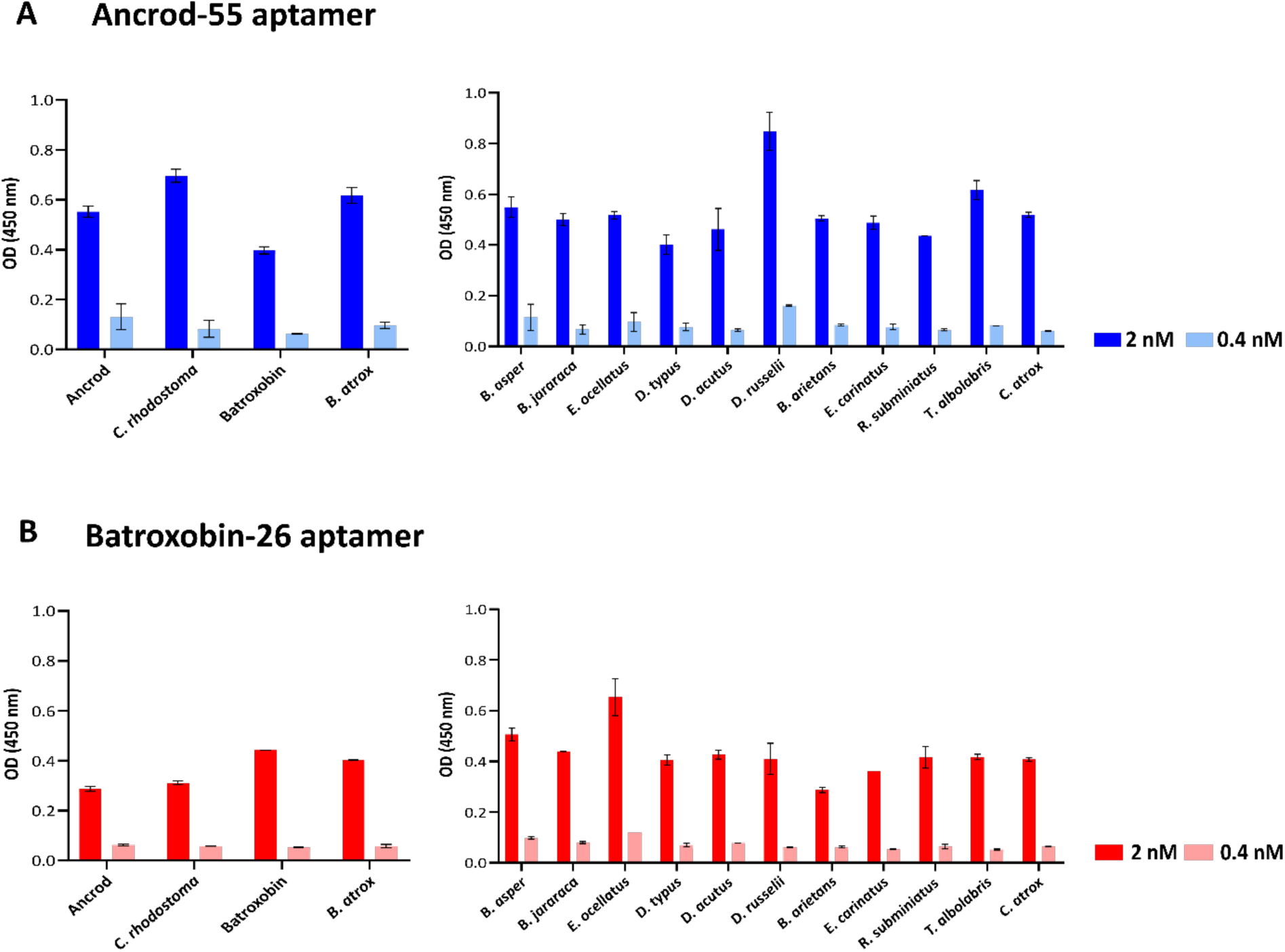
Quantification of binding levels between the selected ssDNA aptamers (Ancrod-55 and Batroxobin-26) and the SVSP recombinant toxins (Ancrod and Batroxobin) and a panel of native snake venoms by ALISA. Data is shown for the aptamers Ancrod-55 **(A)** and Batroxobin-26 **(B)**. The data on the left shows the binding levels obtained against the two recombinant toxins that the aptamers were selected against and the corresponding snake venoms that these toxins are derived from. The data on the right shows binding levels obtained against a broad panel of 11 distinct snake venoms. All data represent the mean of duplicates measured at 450 nm, and error bars represent the SD of the duplicate measurements.

Next, the ALISA was used to explore the capability of each aptamer to bind to a panel of 11 distinct venoms that exhibit considerable variation in toxin composition and were sourced from taxonomically diverse snake species (81). The binding patterns obtained from these experiments suggest that the two aptamers, each selected against a single SVSP toxin, recognise distinct snake venoms to a similar extent as the toxins and homologous snake venoms described above. Thus, at the 2 nM concentration tested, the aptamer Ancrod-55 exhibited a mean optical density (OD) (450 nm) of 0.53 nm (range 0.40-0.84 nm) against the 11 venoms, compared with 0.55 nm and 0.40 nm against Ancrod and Batroxobin, and 0.70 nm and 0.62 nm against *C. rhodostoma* and *B. atrox* venom, respectively (Figure 3A). Similarly, Batroxobin-26 exhibited a mean OD of 0.43 nm (range 0.29-0.65 nm) against the various venoms, compared with 0.29 nm and 0.44 nm against Ancrod and Batroxobin, and 0.31 nm and 0.40 nm against *C. rhodostoma* and *B. atrox* venom, respectively (Figure 3B). While the range in binding levels observed suggests a considerable degree of variation in terms of venom recognition, it is possible that this might simply reflect the relative abundance of SVSP toxins present in these different venoms.

### Aptamers inhibit toxin-induced fibrinogenolysis and fibrinogen depletion

Many SVSP toxins exert a functional activity akin to thrombin by acting to cleave the α-or/and β- chains of fibrinogen, typically resulting in the release of fibrinopeptides rather than fibrin and thus ultimately contributing to dysregulation of coagulation via depletion of fibrinogen (82, 83). To assess whether the binding exhibited by the highest affinity ssDNA aptamers were capable of inhibiting the functional activities known from SVSPs, we first used degradation SDS-PAGE to explore protection against fibrinogenolysis. Our findings revealed that, in line with the known functional activities of native ancrod and batroxobin and recent work (76) both recombinantly expressed toxins cleaved the α-chain of fibrinogen (Figure 4A and 4B). We then assessed the inhibitory capability of the two candidate ssDNA aptamers at preventing this functional activity. Both Ancrod-55 and Batroxobin-26 aptamers protected against fibrinogenolysis caused by the corresponding recombinant toxin, resulting in visualisation of the α-chain of fibrinogen on the gel (Figure 4A and 4B). We next sought to explore whether these two aptamers could also protect against native venom, which might contain additional related (or unrelated) proteins that could contribute to global fibrinogenolysis. Despite this possibility, our data showed that the aptamer Ancrod-55 inhibited the fibinogenolytic effect of *C. rhodostoma* venom on both the α- and β-chains, while Batroxobin-26 aptamer prevented *B. atrox* venom-induced cleavage of the α-chain of fibrinogen (Figure 4C and 4D). These findings suggest that the binding specificities obtained during aptamer selection translate into inhibitory capabilities.

**Figure 4.**
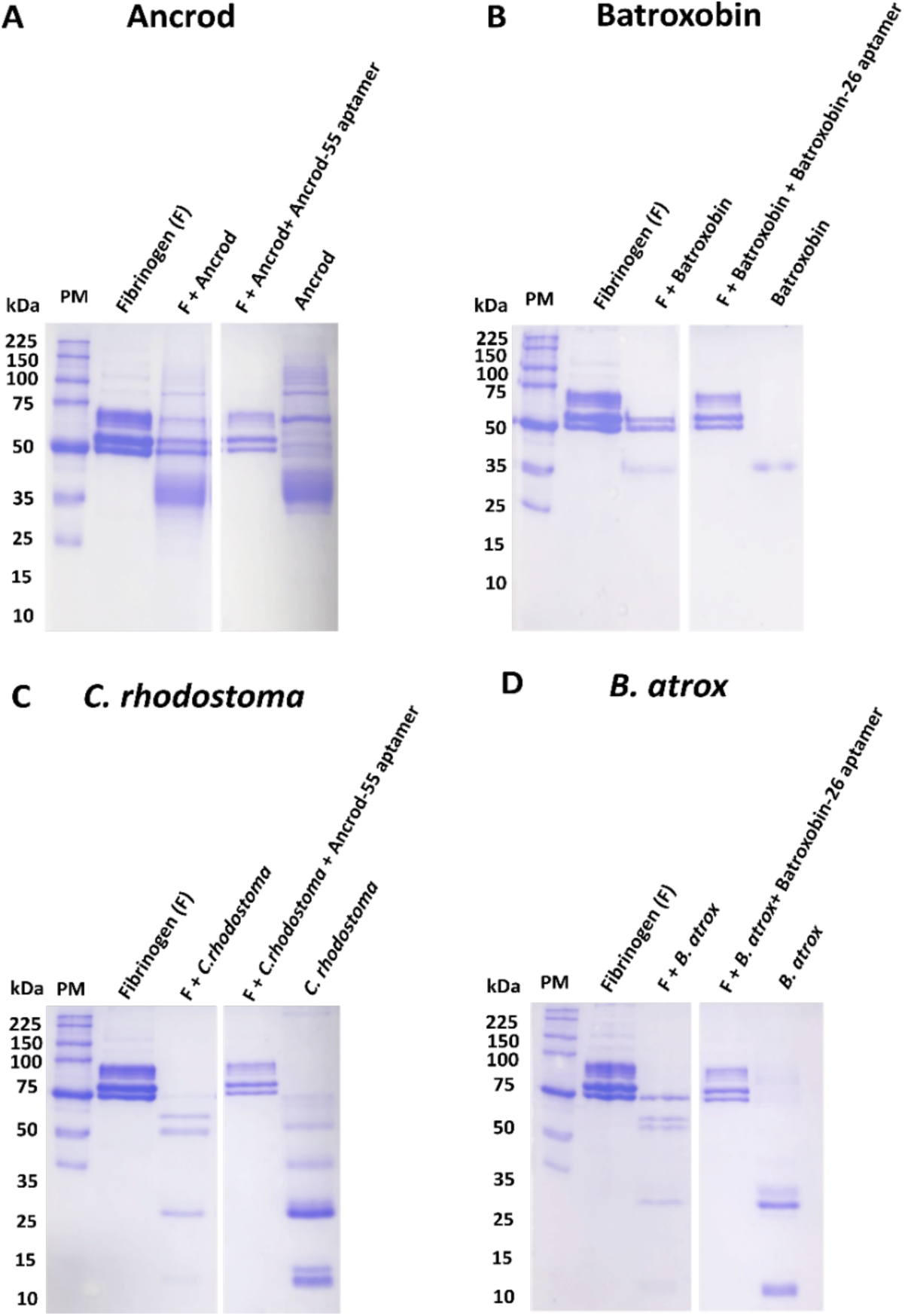
Degradation SDS-PAGE gel electrophoresis demonstrates that ssDNA aptamers directed against Ancrod and Batroxobin inhibit the fibrinogenolytic activity of recombinant toxins and corresponding snake venoms. **(A)** Fibrinogenolytic activity of Ancrod and inhibition by the aptamer Ancrod-55. **(B)** Fibrinogenolytic activity of Batroxobin and inhibition by the aptamer Batroxobin-26. **(C)** Fibrinogenolytic activity of *C. rhodostoma* venom and inhibition by Ancrod-55. **(D)** Fibrinogenolytic activity of *B. atrox* venom and inhibition by Batroxobin-26. Figure shows reduced 8% SDS-PAGE visualised by Coomassie blue staining. All panels have the following layout: Lane 1, protein marker (PM); Lane 2, human plasma fibrinogen (3 μg); Lane 3, toxin (Ancrod or Batroxobin, 7 μg) or venom (*C. rhodostoma* or *B. atrox*, 7 μg) + fibrinogen; Lane 4, toxin or venom + fibrinogen + aptamer (Ancrod-55 or Batroxobin-26, 1 pM); Lane 5, toxin or venom only.

The clotting time of diluted plasma with a standard thrombin concentration is inversely related to the fibrinogen concentration (79). To further explore the inhibitory effect of the selected high-affinity aptamers against both SVSP toxins and native snake venom, we measured fibrinogen depletion using native venom spiking experiments with fresh frozen plasma (FFP), platelet-poor plasma (PPP) and cryoprecipitated anti-haemolytic factors (Cryo-AHF). Our results revealed that both Ancrod and Batroxobin, as well as native venom from *C. rhodostoma* and *B. atrox*, stimulate fibrinogen consumption in a broadly comparable manner across the three plasma samples, resulting in reductions versus the thrombin-stimulated saline control (FFP, <1.5 g/L vs 3.54 g/L; PPP, <1.5 g/L vs 3.42 g/L; Cryo-AHF, <1.5 g/L vs 2.26 g/L) (Figure 5). Across all three of the different plasma-derived samples utilised, the aptamers Ancrod-55 and Batroxobin-26 reduced the consumption of fibrinogen induced by the toxins and native venoms towards control levels (Figure 5). In FFP, the aptamers reduced the fibrinogen consumption of the two native venoms to near control levels (3.54 g/L), and though the level of inhibition was reduced against the corresponding toxins, fibrinogen concentrations remained considerably higher than the toxin only controls (2.1-2.4 g/L vs 1.15 g/L) (Figure 5A). However, the aptamers were more effective in reducing the consumption of fibrinogen in both PPP and Cryo-AHF, which may at least be partly due to the depletion of platelets in these samples, and despite the toxins seemingly exhibiting increased potency in the Cryo-AHF sample (Figure 5C). In both cases, however, the two aptamers inhibited the depletion of fibrinogen stimulated by both toxins and native venoms to near control levels (PPP, 2.7-2.9 g/L vs 3.45 g/L g/L; Cryo-AHF, 1.8-2.2 g/L vs 2.25 g/L, respectively) (Figure 5B and 5C). Crucially, irrespective of the plasma sample used, the aptamers showed highly comparable inhibitory potencies with those obtained with gold-standard commercial antivenoms, as evidenced by the resulting equivalent fibrinogen concentrations observed (Figure 5).

**Figure 5.**
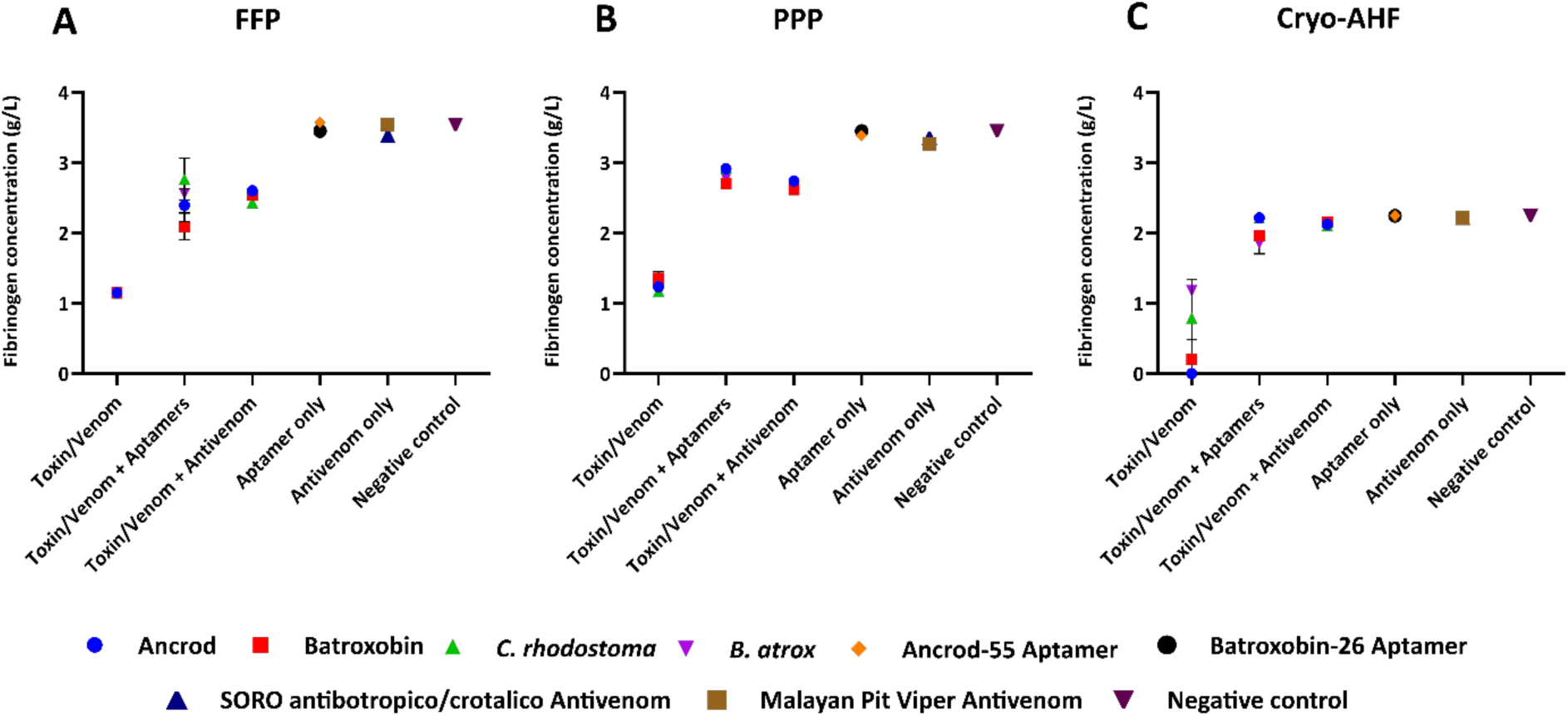
ssDNA aptamers inhibit fibrinogen consumption induced by recombinant toxins and corresponding snake venoms. Fibrinogen concentrations were quantified using an excess of thrombin to convert fibrinogen to fibrin in diluted citrated human plasma samples, and the data shown here represents those obtained from three different types of plasma samples, namely: FFP **(A)**, PPP **(B)** and Cryo-AHF **(C)**. Samples were spiked with 0.6 ng of either recombinant toxin or native venom only (i.e., Ancrod, Batroxobin, *C. rhodostoma* or *B. atrox*); toxin/venom + 1 pM of specific aptamer (Ancrod/*C. rhodostoma* with Ancrod-55, Batroxobin/*B. atrox* with Batroxobin-26); or toxin/venom + 0.5 µg of specific commercial antivenom (Ancrod/*C. rhodostoma* with Malayan pit viper antivenom, Batroxobin/*B. atrox* with SORO antibotropico/crotalico antivenom). Controls consisted of aptamer only samples, antivenom only samples, and saline solution (negative control). Error bars represent the SD of duplicate measurements.

### Aptamers reduce toxin- and venom-induced prolongations of the PT and aPTT

The PT measures the time for citrated plasma to clot and specifically assesses the clotting capability of the extrinsic and common coagulation cascades. A prolonged PT can result from an absence or deficiency of one of the clotting factors X, VII, V, II or fibrinogen. Our recombinant toxin/native venom spiking experiments demonstrated that both Ancrod and Batroxobin, and the corresponding venoms from *C. rhodostoma* and *B. atrox*, result in a prolongation of the PT in PPP compared to the saline control (i.e., >18 seconds vs ∼14 seconds, respectively) (Figure 6A). Noticeably, when co-incubated with the corresponding toxins/native venoms, the aptamers Ancrod-55 and Batroxobin-26 caused a substantial reduction in the PT compared with the toxin/native venom-only samples, resulting in the PT approaching the level of control readings (14.5-15.5 seconds vs ∼14.0 seconds, respectively), and thus demonstrating inhibition of toxin activity (Figure 6A). These reductions were highly comparable, and thus equipotent, to those obtained with commercially available antivenoms (14-16 seconds). Contrastingly, when repeated in the Cryo-AHF samples, which contain the clotting factors XIII, XI, VIII, V, Fibrinogen and vWF, neither of the toxins or snake venoms consistently affected the PT (Figure 6B).

**Figure 6.**
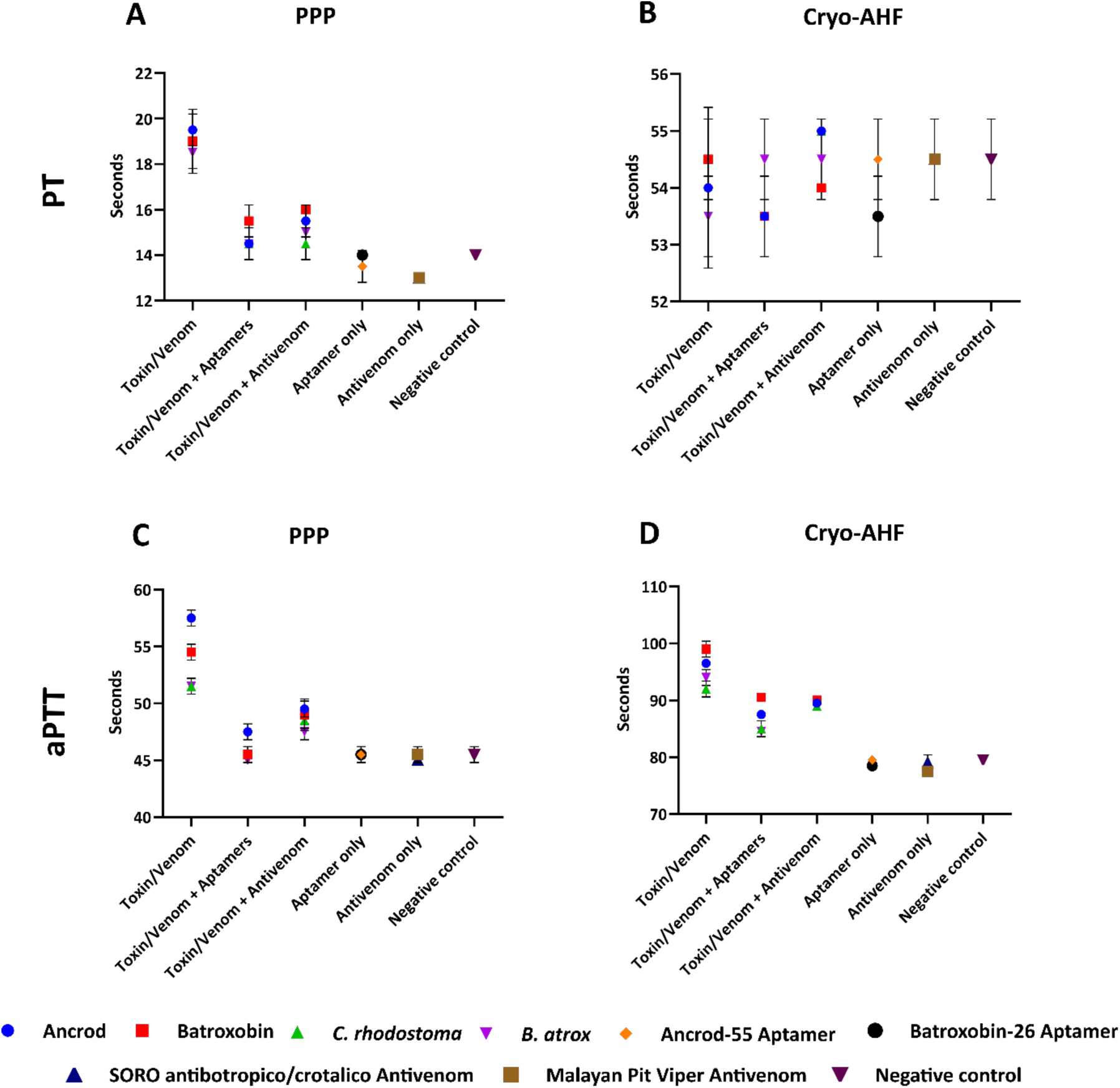
Prolongation of the prothrombin time (PT) and activated partial thromboplastin time (aPTT) stimulated by recombinant toxins and corresponding snake venoms is reduced by the ssDNA aptamers. The PT, which measures the combined effect of clotting factors of the extrinsic and common coagulation pathways (in seconds), and the aPTT, which measures the combined effect of the clotting factors of the intrinsic and common coagulation pathways, were quantified using PPP and Cryo-AHF samples. **A)** PT in PPP, **B)** PT in Cryo-AHF, **C)** aPTT in PPP, and **D)** aPTT in Cryo-AHF. Samples were spiked with 0.6 ng of either recombinant toxin or native venom only (i.e., Ancrod, Batroxobin, *C. rhodostoma* or *B. atrox*); toxin/venom + 1 pM of specific aptamer (Ancrod/*C. rhodostoma* with Ancrod-55, Batroxobin/*B. atrox* with Batroxobin-26); or toxin/venom + 0.5 µg of specific commercial antivenom (Ancrod/*C. rhodostoma* with Malayan pit viper antivenom, Batroxobin/*B. atrox* with SORO antibotropico/crotalico antivenom). Controls consisted of aptamer only samples, antivenom only samples, and saline solution (negative control). Error bars represent the SD of duplicate measurements.

We employed the aPTT in the same manner as the PT, except in this case to measure clotting in the context of the combined effect of the intrinsic and common coagulation pathways (i.e., Factors II, V, VIII, IX, X, XI, XII & fibrinogen). Ancrod, Batroxobin, *C. rhodostoma* venom and *B. atrox* venom all caused a prolongation of the aPTT in both PPP and Cryo-AHF samples when compared with the negative control (Figure 6C and 6D). In both cases, the ssDNA aptamers Ancrod-55 and Batroxobin-26 caused reductions in these prolongations, though with differences in inhibitory potencies observed between PPP and Cryo-AHF. In the case of PPP, the aptamers showed complete inhibition of toxin and native venom activity by reducing the aPTT to control levels, except for the Ancrod and Ancrod-55 aptamer combination which showed substantial inhibition approaching control levels (Ancrod, 57 seconds; Ancrod + Ancrod-55 aptamer, 48 seconds; normal control, 45 seconds) (Figure 6C). Inhibition was also observed for the Cryo-AHF samples, where the two aptamers reduced the aPTT prolongation (84-91 seconds) in all cases when compared with each of the toxin/ venom only measurements (91-100 seconds), though these reductions did not reach control levels (∼80 seconds) (Figure 6D). For all aPTT comparisons, the aptamers slightly outperformed the levels of neutralisation obtained with the commercial antivenoms, except for inhibition of Ancrod in the Cryo-AHF samples, where equipotency was observed. These findings are noteworthy when considering the differences in doses of these therapeutics (aptamer, 1 pM; antivenom 4.55 pM).

## Discussion

Though current antivenom treatments are life-saving therapeutics, they possess several limitations that restrict their clinical efficacy and utility for tackling tropical snakebite. Consequently, a variety of novel experimental approaches are currently being employed to develop new therapeutic interventions that circumvent these existing limitations with snakebite treatment. Next-generation antivenoms consisting of monoclonal antibodies (mAbs) selected for their desirable specificities show much promise in this regard, particularly since different antibody formats can be adapted to specific pharmacodynamic and pharmacokinetic needs (84). However, to date, little effort has been devoted to exploring antibody alternatives despite outstanding concerns relating to the potential cost of mAb-based snakebite treatments. In other fields, ssDNA aptamers have emerged as promising alternatives to antibodies for both therapeutic application and their use as diagnostic tools (38-40), and several ssDNA aptamers have been successfully selected and applied against a wide range of targets including small molecules (85) and toxins (37, 86-89). Despite this prior research, ssDNA aptamers remain largely unexplored for their potential utility as snakebite therapeutics. Thus, in this proof-of-concept study we sought to select ssDNA aptamers against fibrinogenolytic SVSP toxins found in medically important viper venoms, and to evaluate their potential utility as novel toxin-inhibiting molecules via *in vitro* cross-reactivity and neutralisation studies.

Following fourteen cycles of SELEX selection, we selected two pools of aptamers directed against the recombinantly expressed SVSP toxins Ancrod and Batroxobin, which are found in native form in the venoms of the south-east Asian pitviper *C. rhodostoma* and the south American lancehead pitviper *B. atrox*. Although increases in binding to each target increased steadily over SELEX selection cycles, by round 11 major increases in fluorescence intensity, indicating enrichment in binding to the target, had occurred. Thereafter counter-selection was employed to deplete binders directed towards the target carrier (beads), and two further cycles of enrichment were employed, ultimately resulting in highly comparable fluorescence intensity patterns obtained during the selection process to the two different targets. Sequencing 100 targets from the resulting two pools of ssDNA aptamers yielded thirteen and nine unique aptamers directed against Ancrod and Batroxobin, respectively, and analyses of resulting dissociation constants enabled the selection of the highest affinity aptamers against each, which ultimately exhibited desirable low nanomolar K_d_s (i.e., Ancrod-55 aptamer, 3.0 nM; Batroxobin-26 aptamer, 4.7 nM). These dissociation constants compare favourably with those obtained during ten rounds of diagnostic ssDNA aptamer selection against the neurotoxin β-bungarotoxin from the venom of *B. multicinctus* (>65.9 nm), for example (35).

Although Ancrod and Batroxobin are members of the same gene family (i.e., they are both snake venom serine proteases), they have different molecular weights (26–33 kDa) and share only 67% amino acid sequence similarity (90). Nonetheless, ALISA binding experiments demonstrated that the highest affinity aptamers selected and directed against one of these SVSP toxins, were capable of cross-recognising and binding reciprocally to the other recombinant toxin under study here. Moreover, these experiments also revealed that both of the aptamers (i.e., Ancrod-55 and Batroxobin-26) were capable of binding to native toxins present in snake venoms that corresponded to the source of the recombinant toxins (i.e., *C. rhodostoma* and *B. atrox* venom), and indeed a broader variety of unrelated and distinct native snake venoms. Broadly speaking each of the aptamers exhibited comparable moderate binding levels to each of the venoms tested, presumably via interaction with other SVSP toxins present within them, though Ancrod-55 generally exhibited higher binding levels to these different venoms than Batroxobin-26. However, we cannot rule out that non-specific interactions between the aptamers and other venom toxins might be contributing to the overall binding levels obtained in these experiments.

Both Ancrod and Batroxobin are coagulopathic SVSP toxins capable of stimulating fibrinogen consumption and prolonging clotting times (13, 15, 16). Indeed, most SVSPs are fibrinogenolytic enzymes that result in the depletion of circulating fibrinogen during envenoming. Using a panel of *in vitro* bioassays, we demonstrated that recombinant ancrod and batroxobin used for aptamer selection exhibit functional activities consistent with native toxins, including fibrinogenolytic activity via cleavage of the α-chain of fibrinogen, the reduction of fibrinogen plasma concentrations, and the prolongation of clotting times in PT and aPTT assays. Crucially, the coincubation of the ssDNA aptamers with the corresponding recombinant toxins abolished the cleavage of fibrinogen and reduced the consumption of fibrinogen in human plasma samples towards control levels, demonstrating that the binding observed translates into a degree of functional inhibition. Similar results were observed when measuring the PT and aPTT in toxin-spiked platelet-poor plasma, with the homologous aptamers reducing the prolongation of clotting times stimulated by each toxin to near control levels. Inhibition also extended to native snake venoms, with the aptamers inhibiting their fibrinogenolytic and delayed clotting time effects in a highly comparable manner to those obtained with the recombinant toxins. These findings suggest that SVSP toxins likely dominate the fibrinogenolytic effect of *C. rhodostoma* and *B. atrox* venoms and are being effectively inhibited by the ssDNA aptamers Ancrod-55 and Batroxobin-26, though we cannot rule out some contributory interactions between the aptamers and distinct coagulopathic toxins found in these venoms. Irrespectively, it is worth noting that the toxin inhibitory activities demonstrated here by the selected high-affinity aptamers are potent, with low concentrations (1 pM) inhibiting toxin and venom activities observed across the various assays, which compares very favourably to the doses required by the conventional polyclonal antivenoms to effect comparable neutralisation (4.55 pM). Future work should explore whether the *in vitro* inhibitory data obtained here might translate into *in vivo* protection against venom-induced toxicity in preclinical animal models, or whether toxin-family directed aptamers, such as those identified here, need to be combined with other inhibitory molecules (i.e., other aptamers or different inhibitory molecules like mAbs or small molecule drugs) targeting distinct toxin families to prevent severe envenoming pathology *in vivo*.

Nonetheless, the proof-of-concept findings described here reveal that rationally selected ssDNA aptamers hold great value for exploration as novel inhibitory molecules capable of broadly inhibiting snake venom toxins, such as SVSPs. However, the potential therapeutic utility of aptamers does not come without a number of challenges. One major limitation is that ssDNA aptamers are susceptible to nucleases and this, combined with their small size, results in short half-lives *in vivo*, which can be as low as two minutes due to DNA degradation and rapid clearance (49). However, aptamers also possess great flexibility and, as such, can be modified to increase their stability and their resulting half-life, with conjugation to a higher molecular weight protein carrier, for example, liposomes, cholesterol or PEG, previously been demonstrated to successfully decrease the clearance rate from plasma to acceptable levels (57, 59, 91). Other limitations include the initial selection process often being labour intensive and time-consuming, though downstream production of the resulting identified ssDNA aptamers offers a number of advantages over mAb production, for example, due to their size and chemical synthesis. Finally, to date, it is worth re-emphasising that only a single aptamer has passed through an FDA-approval process (Macugen® [Pegaptanib]) (62), and as such regulatory challenges are likely to remain for exploiting ssDNA aptamers as new therapeutics.

Despite these limitations, aptamers possess several clear advantages in terms of therapeutic characteristics over the polyclonal antibodies currently used for treating snakebite. For example, aptamers are chemically synthesised and thus do not require the use of ethically and financially costly experimental animals, and the resulting product exhibits a relative lack of batch to batch variation, unlike biologic antivenom (92). Further, aptamers exhibit desirable stability and are likely more resistant to harsh conditions than antibodies (93), which may be of particular benefit in cold-chain unstable parts of the tropics that suffer a high burden of snakebite. Finally, the large-scale cost of manufacturing aptamers seems likely to be relatively inexpensive compared with existing polyclonal antivenoms. Currently, manufacturing costs for aptamers range from USD 140-280 per dose (91), and while the dose efficacy of aptamers in the context of snakebite remains to be established, it seems likely that aptamer-based treatments could reduce the high costs currently borne by tropical snakebite victims, which often exceed USD 1,000 in sub-Saharan Africa, for example (31). Although much work needs to be undertaken before such benefits can be realised, the proof-of-concept data described herein add further weight to the potential utility of aptamers as future snakebite treatments and strongly advocate for their continued exploration as a novel therapeutic modality for inhibiting pathogenic snake venom toxins.

## Acknowledgements

The authors wish to thank Paul Rowley and Edouard Crittenden for expert snake husbandry at LSTM. N.A. acknowledges support a PhD scholarship provided by the Ministry of Higher Education, Saudi Arabia and the Royal Embassy of Saudi Arabia, Cultural Bureau, London. N.R.C. acknowledges support from a Sir Henry Dale Fellowship [200517/Z/16/Z] jointly funded by the Wellcome Trust and Royal Society. This research was funded in part by the Wellcome Trust [200517/Z/16/Z]. For the purpose of open access, the author has applied a CC BY public copyright licence to any author accepted manuscript version arising from this submission.

## Author Contributions

N.A., N.R.C. and M.Z. designed the study. N.A. and N.R.C. chose and provided the analyte. R.C. planned the SELEX experiments. N.A. and R.C. carried out the SELEX and the DNA sequencing experiments. J.A. planned the ALISA experiments. N.A. performed all other experiments and analysed the data. N.A. wrote the manuscript with input from N.R.C. All authors reviewed and edited the final version of the manuscript.

